# DIFFERENTIAL SCANNING CALORIMETRY: A SUITABLE METHODOLOGY TO PROBLEM THE BINDING PROPERTIES OF BCL-2 INHIBITORS

**DOI:** 10.1101/2023.10.07.561353

**Authors:** Bircan Dinc, Berna Dogan, Thomas Mavromoustakos, Serdar Durdagi

## Abstract

The Differential Scanning Calorimeter (DSC) technique is commonly used for quantitative analysis of protein stability, and it is one of the methods used to quantify protein-ligand interactions. DSC provides denaturation curves that can be a source of data on binding constants. B-cell leukemia/lymphoma-2 (BCL-2) is an antiapoptotic protein, one of the overexpressed proteins in cancer cells. Here, binding constants for BCL-2 of nine BCL-2 inhibitors were evaluated by utilization of DSC measurements. The binding free energies of suggested inhibitors of BCL-2 were measured by melting temperature, T_m,_ and heat capacity, Cp. Evaluated BCL-2 binding parameters of candidate molecules agreed with our previously reported biological assays and molecular simulations indicating that the DSC method could be applied for the determination of free energies of enzyme inhibitors with high accuracy. Therefore, our results showed that this fast and relatively cheap technique can be conducted in high throughput drug screening analyses to compare the binding affinity of a set of compounds against a specific target protein. BCL-2 and most of its inhibitors have hydrophobic cavities, and a definite percent of dimethyl sulfoxide (DMSO) was advised to increase the solubility. Our work also suggests that the presence of the recommended high concentration of DMSO as the solvent significantly changes the affinity of the inhibitors to BCL-2, revealing similar DSC curves for all inhibitors.

## Introduction

There is a need for advanced screening methods that can rapidly and accurately evaluate target protein-inhibitor binding properties, along with targeted drug design studies (i.e. molecular docking) and *in vitro* biological experimental assays [1]. These screening methods can have a significant impact in determining the success of new therapeutics and in early preclinical drug research. When the ligand binds to the free protein, there is an increase in the melting temperature of the ligand-bound protein as a function of increasing ligand concentration. Depending on the strength of the binding interaction, marked changes in the thermogram of the protein occur in the presence of the ligand. Thermograms produced from differential scanning calorimetry (DSC) are sensitive to protein-ligand interactions. Surprisingly less attention from ITC (isothermal titration calorimetry), DSC can be used as a superior screening alternative with small volume and easy sample preparation, measurements that can be taken in less than 90 minutes without the need for prior binding knowledge. It does not require entering any prior information about the binding activity [2-4].

ITC measurements are also a widely used method for the determination of protein-ligand binding constant and binding stoichiometry. However, DSC has the advantage in situations where binding is very slow, where ligand dissolution occurs in the presence of high concentrations of organic solvents such as DMSO, or where binding is too high an affinity to be measured by ITC [5]. Most of the synthesized ligands have dissolution problems, and it is necessary to add organic solvents such as dimethyl sulfoxide (DMSO) or dimethylformamide (DMF) to the solvent in certain proportions [6]. The presence of these solvents poses a problem in the measurements taken with ITC. The heat generated by protein-ligand binding generates a small heat of attachment. During DSC measurements, it is possible to match the protein and solvent placed in the reference cell, the protein, ligand, and solvent placed in the sample cell, and the organic concentrations in the titrations. In addition, most of the DSC systems used require less volume and mass of protein and ligand than ITC [7, 8].

The degree of stabilization of the ligand-protein complex against thermal denaturation in DSC is compared. Therefore, the ligand is more advantageous than ITC in cases where it binds to the target protein with high affinity. While long-term interaction of protein and ligand is allowed in DSC experiments, this interaction takes place on the order of minutes in ITC experiments. In addition, the ligand can be measured at different concentrations, and how its binding changes depending on temperature and concentration can be examined [8, 9].

Therefore, in the current study, we used a well-known oncotarget B-cell leukemia/lymphoma-2 (BCL-2) as a model target and explored the usage of DSC in the screening of small molecules on this macromolecular target. BCL-2 family proteins tightly regulate the intrinsic apoptosis process and initiate apoptosis by activating caspase-9 [2, 10, 11]. The BCL-2 proteins, which include evolutionarily related proteins, are a family in which all members maintain sequence patterns called the BCL-2 homology (BH) domain. The members of this family are generally divided into three main classes: (i) pro-apoptotic activators, (ii) pro-apoptotic effectors, and (ii) anti-apoptotic regulators. The first class of proteins consists of activator proteins such as BIM, BID, PUMA, containing only the pro-apoptotic BH domain 3 (BH3). Immediately after their activation, they act as molecular guards that connect the outer spurs to the mitochondrial pathway.

[12] The second class BCL-2 proteins include multi-domain protein pro-apoptotic effectors such as BAX, BAK, and each has three BH domains. These proteins initiate caspase activation by disrupting the integrity of the mitochondrial outer membrane leading to the free circulation of cytochrome C into the cytoplasm, resulting in cell death. The most promising and important method to activate and increase the mechanism of apoptosis is to increase the amount of pro-apoptotic activator proteins (BCL-2 members containing only the BH3 region) or to block one of the anti-apoptotic BCL-2 counterparts [4-6, 13]. It has been shown in different studies that anti-apoptotic BCL-2 members are overexpressed in cancer cells [8, 9, 14] and BCL-2 upregulation seems to prevent cells from undergoing apoptosis. It has also been observed that cancer cells show oncogene dependence on anti-apoptotic proteins [15, 16]. For these reasons, the use of BCL-2 proteins as a target in anti-cancer therapy seems to be a promising method.

One of the most important problems during screening studies of molecules is solubility. The use of DMSO at different concentrations as a solvent is quite common in these studies. Studies can be conducted with DMSO ratios reaching up to 70%. It revealed both stabilizing and denaturing effects of DMSO according to the concentration. DMSO is an organic polar solvent with a boiling point of 189 °C. Due to its high boiling temperature, it is preferred because of its stabilizing and solubility-enhancing effects at certain temperatures. In this study, the effect of the organic cosolvents on protein structure and stability was also investigated. It has been revealed that DMSO affects thermal stability but does not affect the unfolding mechanism [17]. However, in some later studies, it was determined that even a concentration of 2.5% affected the unfolding pathway [18]. Thus, in the current study, the effect of the used solvent on the structural stability of the ligand and protein: ligand complexes were also investigated.

In our recently reported study [19] we also proposed seven potent inhibitors (**58** (AJ-292/12931005), **ind-199** (AG-205/12549135), **43** (AO-081/41887762), **243** (AN-698/40780701), **258** (AK-968/12163470), **292** (AK-968/11842328), **ind-435** (AN-329/13484046)) that have been screened via computational approaches from a large molecular library (Specs SC) and confirmed to be hit BCL-2 inhibitors by *in vitro* studies (see Table 2 taken from our previous study).

In this study, we measured the thermodynamic parameters of binding of previously suggested hit inhibitors [19] of BCL-2 anti-apoptotic protein by employing DSC measurements. The solubility features of BCL-2 inhibitors with different solvents and their interaction with BCL-2 were evaluated and interpreted by considering the DSC thermograms, and melting temperature (T_m_) values. In addition, changes in thermodynamic parameters were also determined specifically following thermodynamic data were calculated from DSC measurements: change in enthalpy (ΔH), entropy (ΔS), and Gibbs free energy (ΔG) due to ligand binding. These evaluated values for each hit inhibitor molecule have been compared with the computational and *in vitro* results obtained in our previous studies to evaluate the performance of DSC measurements for protein-ligand interactions.

## Materials and Methods

### Effect of Solvents on the Structural Stability

The product data sheet of known BCL-2 inhibitors (i.e., Navitoclax and Venetoclax), suggests solvents containing different concentrations of DMSO for *in vitro* and *in vivo* studies. Navitoclax and Venetoclax are sold as powder samples and stock solutions in 100% DMSO. Therefore, measurements were performed in 10% and 100% DMSO, as well as a 3rd solvent recommended for in vivo experiments [20].

#### Target Protein and Inhibitors

BCL-2 active humans (recombinant, expressed in *E. coli*) were purchased from Sigma. Navitoclax, Venetoclax, Sulfobutylether-β-Cyclodextrin (SBE-β-CD) were purchased from MedChemExpress. BCL-2 inhibitors were purchased from SPECS [27]. 0.9 M NaCl was dissolved in 100 ml distilled water to prepare 0.9% saline solution. 2 g of SBE-β-CD in 0.9% saline was dissolved in 0.9 % saline to make 10 ml with a 20% (w/v) concentration (20% SBE-β-CD in saline). The first solvent was pure DMSO (S1) (Sigma, ≥ 99.9%), the second solvent (S2) is 10% DMSO recommended in all MedChemExpress recommended solvent mixtures for *in vivo* studies. The inhibitor solvent (S3) was 10% DMSO, 90% (20% SBE-β-CD in saline) which was advised for *in vivo* studies for the compounds that have solubility problems [20]. The solvent for BCL-2 protein (S4) is deionized water (dH_2_O) as recommended on the datasheet. 1 mM stock solutions were prepared from the seven inhibitors (**58** (AJ-292/12931005), **ind-199** (AG-205/12549135), **43** (AO-081/41887762), **243** (AN-698/40780701), **258** (AK-968/12163470), **292** (AK-968/11842328), **ind-435** (AN-329/13484046)); and **Navitoclax** and **Venetoclax**.

#### DSC Measurements

An auto DSC Calorimeter (DSC-60 Plus, Shimadzu) was utilized for the measurements of heat capacities and melting temperature. Sealed alumina pans were used, and the instrument was calibrated using indium, under nitrogen atmosphere (flow rate: 50 ml/min). The calibration interval was between 30-160 °C and the temperature rate was 1 °C /min. Thermal Analysis 60 WS data analysis software supplied by Shimadzu Corporation used in calibration and sample measurements. In all measurements, the same amount of buffer was placed in the reference cell as the sample cell.

#### Thermodynamics Property Calculations

Data from DSC measurements are obtained as temperature (T), time (t), and heat flow (q). Heat capacity (Cp) can be calculated using the following equation:

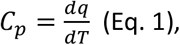

The measured heat capacity difference (Δ*C*_*p*_) between sample pan and reference pan is given by Eq. 2:

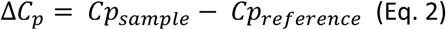

Here, the graph was resketched using Originlab 8.1 software with temperature (°C) on the x-axis and heat flow (mW) on the y-axis. To calculate the specific heat capacity, the heat flow values (mW= mJ/s) were multiplied by the time (s) values. These values were then divided by weight and temperature values. Here, the weight, in mg, was taken as the total value of the ligand contained in the sample pan (Eq. 3):

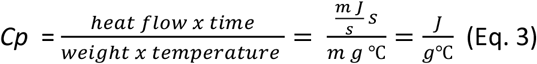

The experimentally derived enthalpy change (ΔH) at a given temperature interval is as follows:

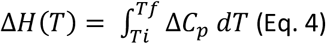

To calculate the ΔH value, first, the heat flow value obtained from the DSC data was divided by the weight: The area under the curve is calculated by integrating the trapezoidal rule and the peak value was determined. Before this calculation, baseline correction has been conducted. The temperature values at the starting and end points of the curve were used to determine the boundaries of the integral. The unit of the calculated area for the enthalpy change was J/g which is converted to kcal/mol.

The experimentally derived entropy (ΔS) shows the free energy association to characterize the thermodynamics of a binding reaction means to determine ΔG. ΔS at a given temperature interval as follows:

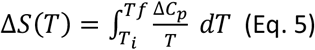

The change of entropy values was obtained by integrating the specific heat by temperature (J/g°C) values obtained using the trapezoid rule (Eq. 5). The binding free energy (Δ*G)* was then determined using Gibb’s free energy relationship:

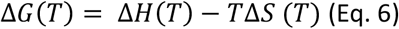

#### Fourier transform infrared spectroscopy (FTIR) Measurements

FTIR measurements were performed with Shimadzu IRAffinity-1S by taking percent transmittance measurements in the wavenumber range of 400-4000 cm^-1^.

#### Statistical analysis

All assays were conducted in triplicate. Mean and standard deviation was estimated using analysis of variance. Data processing and analyses were performed using OriginPro2016 software (OriginLab Corporation, Northampton, MA).

## Results and Discussion

Changes in intramolecular and intermolecular interactions and dynamics of both protein and ligand occur because of their interaction. The binding energy of the protein-ligand complex is comparable to the stability of the ligand-free protein. DSC is a convenient method to study the thermodynamic interactions controlling these conformational changes [21]. In DSC measurements, the degree of stabilization of the bound ligand-protein complex against thermal denaturation is compared. Therefore, DSC is more advantageous than ITC when the ligand binds to the target protein with high affinity. While long-term interaction of protein and ligand is allowed in DSC experiments, this interaction takes place on the order of minutes in ITC experiments. In addition, the ligand can be measured at different concentrations, and how its binding changes depending on temperature and concentration can be examined [22, 23].

DSC measurements allow the determination of heat capacities. It provides the thermodynamics information of the biomolecules such as Gibbs free energy, enthalpy, and entropy in a straightforward manner that enables a deep understanding of the structure-function relationship in biomolecules such as the folding/unfolding of protein and DNA, and ligand bindings. DSC helps with the ranking of binding affinities of ligands against a specific target protein more directly [24].

In the current study, we used the BCL2 target as a model system and screened known BCL2 ligands from our previously published BCL2 inhibitors and FDA-approved BCL2 inhibitor drugs using DSC [25]. Experiments were performed by dissolving the known BCL-2 inhibitors Venetoclax and Navitoclax in different solvents. Recommended DMSO concentrations at the MCE for *in vitro* experiments with Navitoclax and Venetoclax were used [20]. The drugs did not dissolve immediately when DMSO was used, thus they dissolved in a sonic bath for 10 minutes. However, when these two drugs were dissolved in DMSO (1M), they gave similar T_m_ values as DMSO (Figure 1-B).

**Figure 1:**
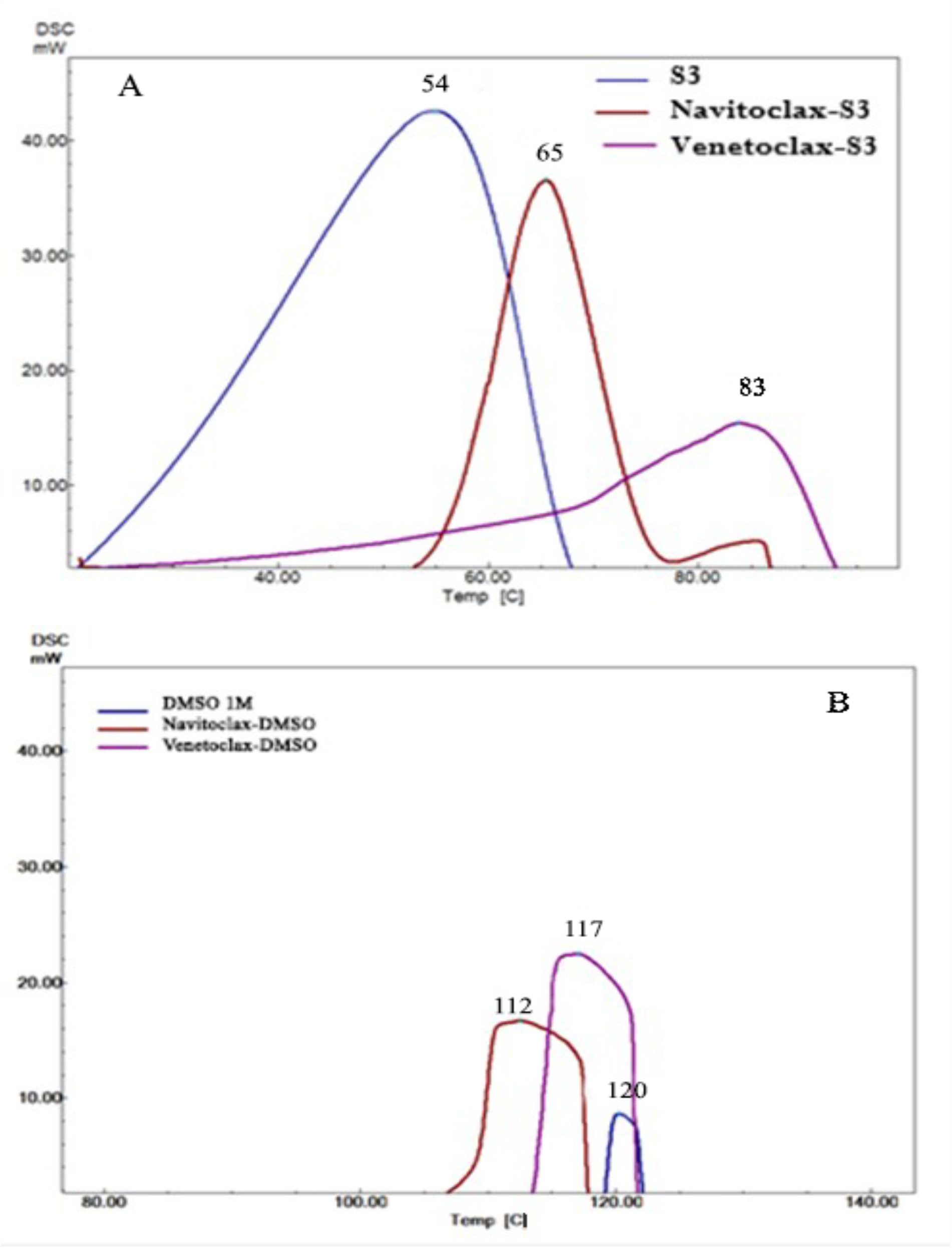
A-DSC thermograms of Navitoclax and Venetoclax solved in S3 (10% DMSO - 90% (20% SBE-β-CD in saline)), B-DSC thermograms of Navitoclax and Venetoclax solved in DMSO (1 M)

Known BCL-2 inhibitors are hydrophobic molecules, to increase their solubilities in water, DMSO is added in certain proportions into the solvent. DMSO can dissolve both polar and non-polar compounds. It is an important polar aprotic solvent miscible with a wide variety of organic solvents and water. Its boiling point is 189 °C which improves the accuracy of test compound concentrations by reducing room temperature evaporation. If the DMSO volume concentration exceeds 10%, the proteins are thought to be unfolded and induce cell death. Therefore, ligand affinity is assessed in the presence of less than 10% of DMSO [26, 27]. There is not much literature on the effects of DMSO on proteins during high-throughput screening. Most of the studies have been carried out at high DMSO concentrations, reaching up to 70%. In these studies, it is explained that DMSO exhibits both stabilizer and denaturation effects [22, 23]. Yang et al. revealed that even 2.5% concentration of DMSO causes a significant change in the unfolding pathway of dimeric bacterial NAD+ synthetase [24]. Dissolving 10% DMSO - 90% (20% SBE-β-CD in saline), Venetoclax, Navitoclax, and other inhibitors in MCE (MedChemExpress)’s recommended solvent for *in vivo* experiments revealed different DSC curves than 1M DMSO (Figure 1). It has been observed that drugs dissolved in 1M DMSO that do not show DSC curves for their folding properties when dissolved in the solvent 3 (S3) containing 10% DMSO - 90% (20% SBE-β-CD in saline), reveal Tm values ranging from 50 to 84 °C (Figure 1-A). It was also confirmed by FTIR results that Venetoclax and Navitoclax dissolved in 1 M DMSO revealed similar spectra (Figure 2) to DMSO. In this case, dissolving the inhibitors in DMSO alters the structural integrity of the inhibitor. The experimental IR of 1M DMSO, 10 % DMSO, S3, and Navitoclax and Venetoclax dissolved in 1M DMSO, S3 shows transmittance values above 3300 cm^-1^ (3482, 3323, 3366, 3482, 3395, 3361 and 3395 cm^-1^ respectively). The transmittance at 3395, 3482 cm^-1^ are attributed to asymmetrically hydrogen bonded water, the transmittance at 3323 cm^-1^ is due to water molecules not participating in hydrogen bonding interactions [28]. The peaks at 3361 and 3366 cm^-1^ may be due to stretching vibrations of N–H [29]. DMSO has asymmetric and symmetric CH stretching modes at 2998, 2938, 2914, and 2878 cm^-1^ [30]. S3 contains 10% DMSO, so the characteristic DMSO peak in the range of 1036-1050 cm^-1^ due to S=O stretching vibrations [31] appeared in the entire FTIR spectra. It is also seen that the peak is more pronounced in those containing 1M DMSO than in those that dissolve in S3. If the fingerprint region of S=O vibrations between 1000-500 cm^-1^ is [32] examined, it is noteworthy that Venetoclax and Navitoclax dissolved in DMSO and DMSO have similar peaks (Figure 2). When the FTIR spectrum of powder Venetoclax was examined, it was detected there were similar peaks to Venetoclax dissolved in S3 in the 1600-600 cm^-1^ range and the orientation of the whole spectrum curve was also similar [33].

**Figure 2:**
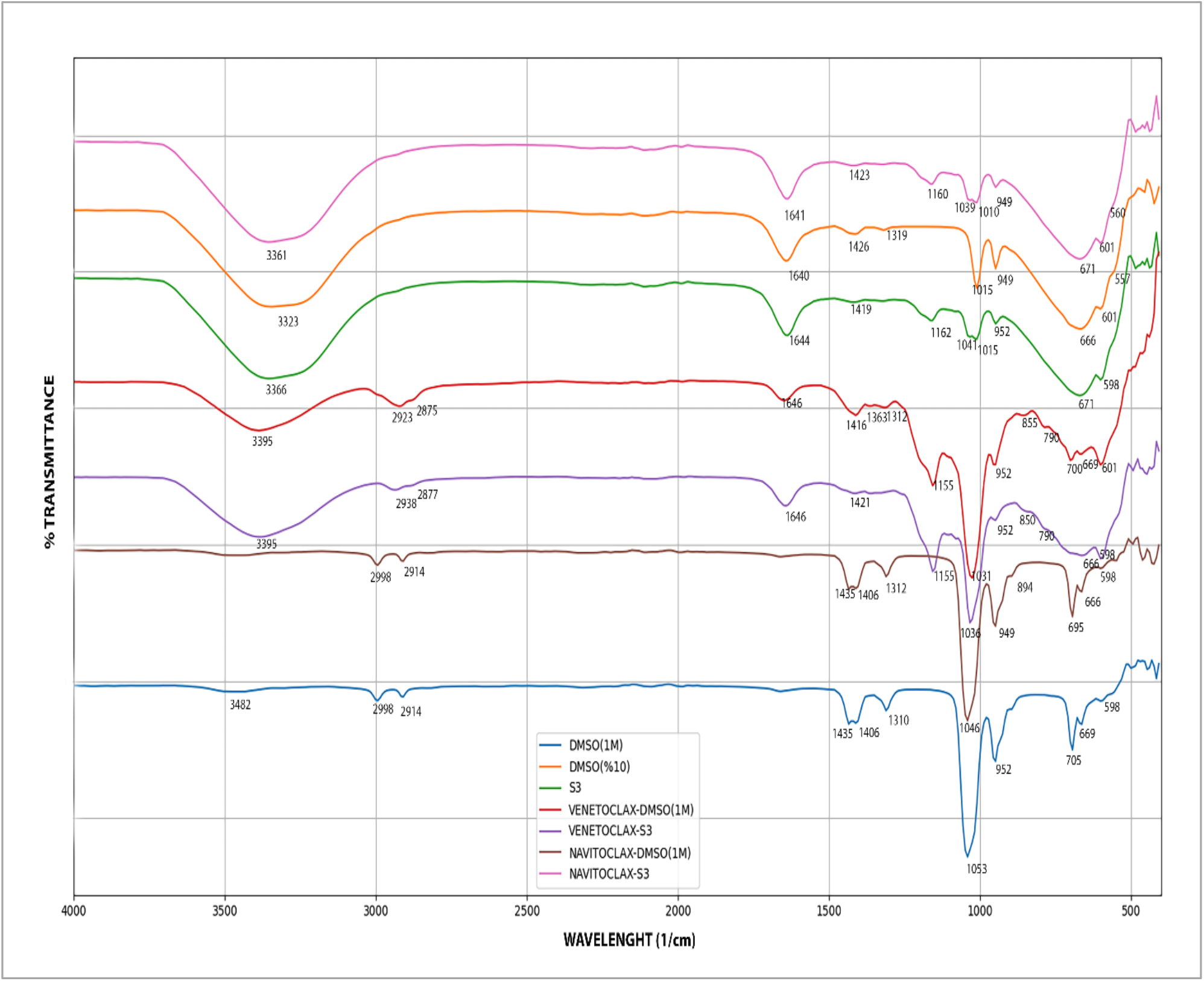
FTIR spectra of DMSO (1 M), DMSO 10%, S3 and Navitoclax, and Venetoclax solved in DMSO and S3 (10% DMSO - 90% (20% SBE-β-CD in saline))

**Figure 3:**
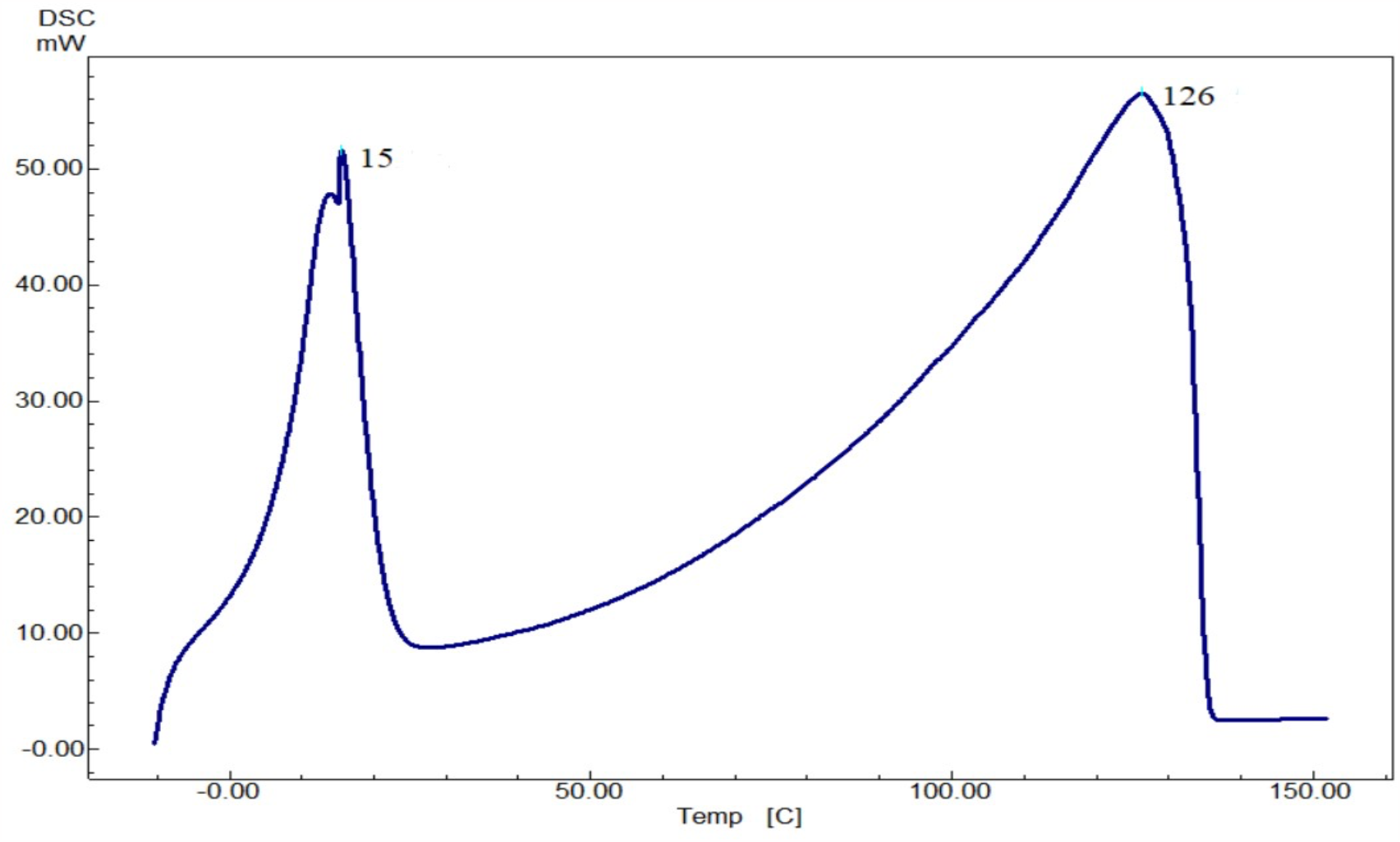
DSC curve of DMSO (1 M) from -10 °C to 150 °C

In previous studies, it was determined that the affinity of 1-adamantane carboxylic acid (ADA) and β-cyclodextrin was significantly reduced in the presence of 5% DMSO. In the experiments, when 5% DMSO and 80% DMSO were used while making measurements in Isothermal Titration Calorimetry (ITC), reproducible results on cyclodextrin derivatives could not be obtained [26]. Cyclodextrins are widely used in commercial formulations to encapsulate hydrophobic drugs. Similar to drug screening techniques, the manufacture of cyclodextrin-based formulations requires the use of a co-solvent to form the hydrophobic drug-cyclodextrin complex [34]. In the screening study here, using 10% DMSO with cyclodextrin (S3) as the solvent yielded much more plausible results for protein-inhibitor interaction.

After the inhibitors were dissolved in 1 M DMSO and S3, they were allowed to interact for 24 hours at +4 °C. The inhibitors dissolved in 1 M DMSO gave unstable peaks at T_m_ temperatures and different DSC thermograms were revealed in each measurement. When S3 was used as the solvent and the measurements were repeated, it was found that each inhibitor produced much more reasonable T_m_ values and DSC thermograms for interactions. They also revealed similar thermogram and T_m_ values in each repeating measurement. Therefore, the solvent containing cyclodextrin and DMSO was used as the solvent of both Venetoclax and Navitoclax and seven other inhibitors.

Peaks appeared at 15 ± 1.6 °C and 126 ± 5 °C in the DSC measurement (heating rate 5 °C /min) results of pure DMSO from -10 °C to 150 °C. In the previous DSC measurements for DMSO, the melting point temperature increased as the DMSO ratio increased in DMSO-water mixtures. Pure DMSO has a melting temperature below 20 °C. In the study here, the heating rate is the same as we performed, but the measurement was started from -80 °C [35]. The known freezing point of DMSO is 19 °C. Depending on the measuring device and conditions, there may be a difference of up to 3 degrees in T_m_ values. In DSC measurements, as the heating rate is increased, the measured peak temperatures increase [36]. In another measurement with the heating rate set to 10 °C /min, DMSO peaked before 150 °C [37]. Molecules dissolved in DMSO produced results around 126 ± 5 °C in our measurements. This confirms the degradation in the structures of molecules. The molecules did not reveal a unique DSC curve (Figure 2-B).

Since the DMSO effect on the structures of the molecules is known, the experiments were continued with the solvent (S3) with a DMSO ratio of 10% and recommended for Navitoclax and Venetoclax. The values resulting from the interaction of each molecule with BCL-2 are shown in figure 4. It was determined that there were different and compatible values according to the in vitro test results (figure 5). The thermodynamics of protein-ligand interactions with DSC can be measured. Results from DSC assays may differ from in vitro experiments because in vitro assays measure the functional inhibition of the target enzyme or protein. The stability of the protein-ligand complex is measured in DSC, which may not always accurately reflect the physiological conditions in which the inhibitor acts. The protein-ligand complex may be more stable in the DSC assay than in the cellular environment, resulting in different DSC and in vitro test results.

**Figure 4:**
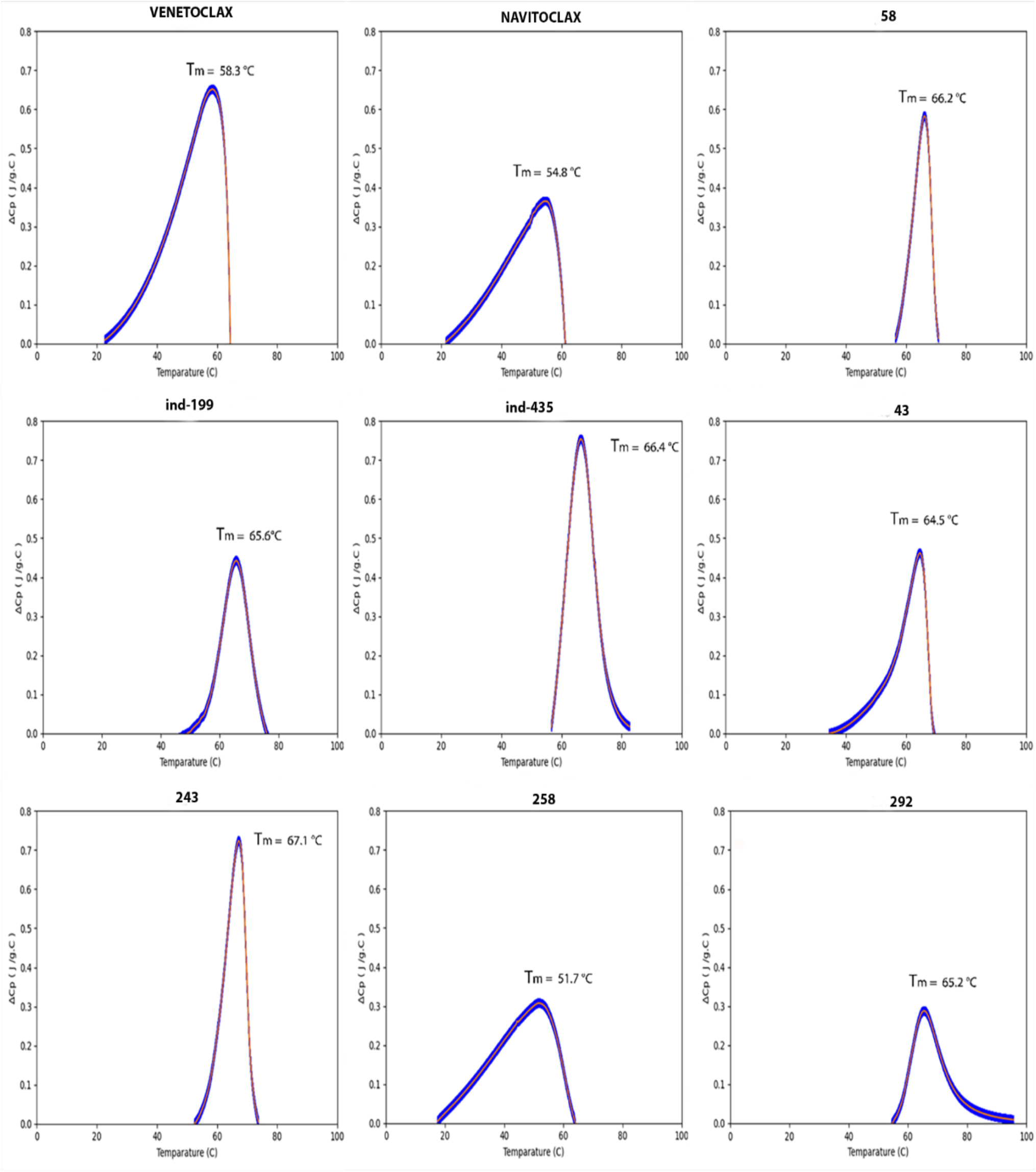
DSC Thermograms of inhibitors. The DSC curves in this figure were obtained by dissolving the compounds in solvent S3.

**Figure 5:**
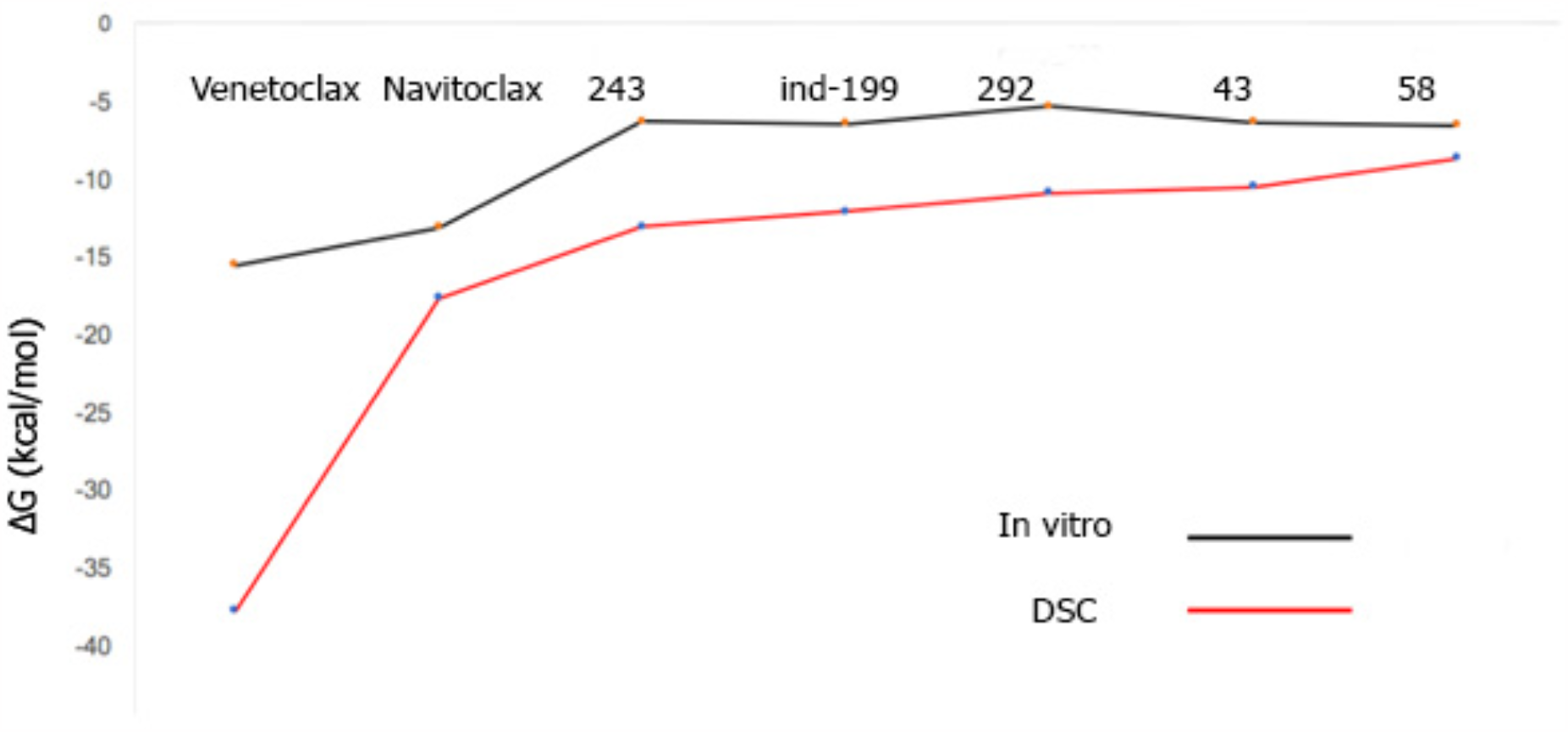
ΔG comparison of DSC and *in vitro* results [25]

DSC measures the stability of the protein-ligand complex; this may not always reflect the physiological conditions in which the inhibitor acts. The protein-ligand complex may be more stable in the DSC assay than in the cellular environment, resulting in differences in values [38].

Navitoclax, Venetoclax, and seven molecules were suggested as new scaffolds for the inhibition of BCL-2 in our previous study and were evaluated by DSC curves (Figure 4). The IC_50_ values of Navitoclax and Venetoclax are known as 0.5 nM (K_i_) and 0.01 nM (K_i_), respectively [39]. The highest binding affinities belong to these two drugs and **ind-435** in DSC measurements. Afterwards, the high affinity measurement results belonged to compounds **ind-435, 243, ind-199, 43, 58** and **292**, respectively (Table 1).

**Table 1:** Thermodynamics parameters of BCL-2 inhibitors (37 °C).

| Inhibitor | T<sub>m</sub> (°C) | ΔH (kcal/mol) | ΔG (kcal/mol) |
| --- | --- | --- | --- |
| Venetoclax | 58.3 | -22.00 | -37.88 |
| **ind-435** | 66.4 | -18.10 | -22.68 |
| Navitoclax | 54.8 | -11.50 | -17.73 |
| **243** | 67.1 | -12.00 | -13.14 |
| **ind-199** | 65.6 | -10.30 | -12.19 |
| **292** | 65.4 | -8.49 | -10.99 |
| **43** | 64.6 | -13.10 | -10.61 |
| **58** | 66.1 | -7.60 | -8.75 |
| **258** | 45.4 | -21.90 | -5.62 |
| **BCL-2** | 57.7 |  |  |

**Table 2.**
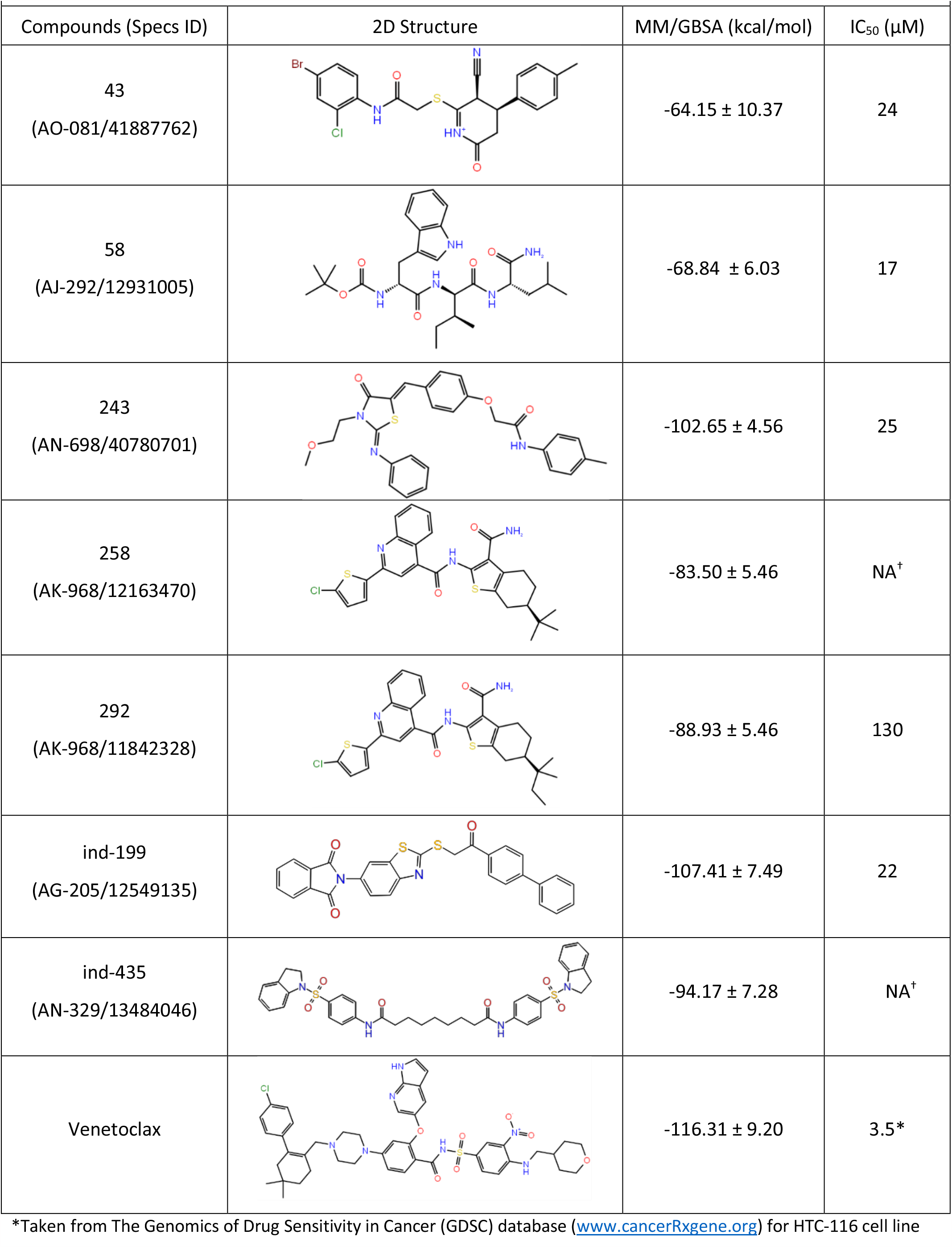
The Specs ID, 2D structure, average MM/GBSA values and IC50 values of compounds.

|  |  |  |  |
| --- | --- | --- | --- |
| Compounds (Specs ID) | 2D Structure | MM/GBSA (kcal/mol) | IC<sub>50</sub> (μM) |
| 43<br>(AO-081/41887762) |  | -64.15 ± 10.37 | 24 |
| 58<br>(AJ-292/12931005) |  | -68.84 ± 6.03 | 17 |
| 243<br>(AN-698/40780701) |  | -102.65 ± 4.56 | 25 |
| 258<br>(AK-968/12163470) |  | -83.50 ± 5.46 | NA<sup>†</sup> |
| 292<br>(AK-968/11842328) |  | -88.93 ± 5.46 | 130 |
| ind-199<br>(AG-205/12549135) |  | -107.41 ± 7.49 | 22 |
| ind-435<br>(AN-329/13484046) |  | -94.17 ± 7.28 | NA<sup>†</sup> |
| Venetoclax |  | -116.31 ± 9.20 | 3.5* |

\*Taken from The Genomics of Drug Sensitivity in Cancer (GDSC) database (www.cancerRxgene.org) for HTC-116 cell line

When a ligand binds to the native state of a protein, the denaturation temperature of the protein and ligand binding will be higher compared to the temperature at which the free protein is denatured (Table 1). The binding between a flexible protein and a small ligand can be complicated by the presence of hydrophobic portions on the surface where it attaches to the protein. The protein-ligand complex can interact with these hydrophobic parts and transfer the bound water in the structure to the solvent. Since all free energy must be reduced in the binding state, the effects of solvent should be considered in protein-ligand binding, as well as energy changes due to non-covalent interactions [40].

In practice, the shape of the protein/ligand endotherm may be quite complex due to the presence of intermediates and/or denatured protein, the sensitivity of the complex to pH and ionic strength, or preferentially binding and stabilizing only a portion of the ligand [40-42]. For many applications, only an estimate of the T_m_ of the free protein versus the Tm of the complex (the temperature at which half the protein/ligand complex molecules are folded, and half unfolded) is required; this information can usually be obtained by visual inspection of endotherms. However, precautions should be taken not to over-interpret the data, as the free energy of complex formation results from the delicate balance between large positive and negative entropic and enthalpic contributions. For example, binding driven by hydrophobic interactions (an entropic effect) tends to result in greater shifts in Tm than enthalpy-directed binding (eg, changes in dissolution). Thus, a large Tm shift observed is not necessarily indicative of high affinity binding, as a number of different affinities with different entropic and enthalpic contributions can result in the same Tm [43].

In the current study our focus is on DSC analysis of BCL-2 inhibitors, determining thermodynamic stability and structural integrity, evaluating drug/ligand binding properties by considering energy due to ligand binding. The thermodynamic binding coefficient for the binding of biologically active ligands to proteins is usually determined using isothermal titration calorimetry. However, this method does not allow to study for very high binding constants [44]. The DSC method is generally used in protein denaturation studies. Protein-ligand interaction can be evaluated according to the differences in the denaturation peak. Calculations can be performed in several steps, yielding quantitative ligand binding results for known protein-ligand concentrations.

The effect of DMSO on protein-ligand interactions may depend on factors such as DMSO concentration, the nature of the protein and ligand, and the specific binding site. In general, lower DMSO concentrations (up to 5-10%) may have minimal effect on the affinity of the interaction, while higher DMSO concentrations (above 10-20%) may cause significant changes in affinity.

At low concentrations, DMSO can increase the solubility of the ligand, making it easier to form a protein-ligand complex [45]. High concentrations of DMSO can disrupt the structure and stability of the protein, resulting in decreased affinity for the ligand. This is because DMSO can act as a denaturant, disrupting hydrogen bonds and other non-covalent interactions critical for protein folding and stability.

Here, in the screening study, the effect of DMSO was also evaluated. In previous studies, it has been determined that DMSO increases beta-sheet formation in proteins, affects the DNA topology and cellular lipid content of cells [46]. However, DMSO is still used as a solvent for most cancer drugs. Considering all these, the effects of DMSO and its ability to inhibit the inhibitor molecule should not be ignored, especially in studies of poorly water-soluble ligands.

## Conclusion

DSC is a method that provides information about binding affinity, stoichiometry and thermodynamics, enthalpy and entropic contributions while measuring protein-ligand interactions. With a small amount of sample it is possible to measure without a prior knowledge input into the system, as in ITC, but experiments should be performed with purified protein and ligand samples. It provides an understanding of the functional effects of protein-ligand interactions rather than their structural details. Depending on the protein-ligand system examined, it can be considered as a highly effective method.

It is important to note that the effect of DMSO on protein-ligand interactions can vary greatly depending on the specific system studied. Therefore, when designing experiments to study protein-ligand interactions, it is important to carefully consider the DMSO concentration and its potential effects.

